# Benchmarking Algorithms for Gene Set Scoring of Single-cell ATAC-seq Data

**DOI:** 10.1101/2023.01.14.524081

**Authors:** Xi Wang, Qiwei Lian, Haoyu Dong, Shuo Xu, Yaru Su, Xiaohui Wu

## Abstract

Gene set scoring (GSS) has been routinely conducted for gene expression analysis of bulk or single-cell RNA-seq data, which helps to decipher single-cell heterogeneity and cell-type-specific variability by incorporating prior knowledge from functional gene sets. Single-cell assay for transposase accessible chromatin using sequencing (scATAC-seq) is a powerful technique for interrogating single-cell chromatin-based gene regulation, and genes or gene sets with dynamic regulatory potentials can be regarded as cell-type specific markers as if in scRNA-seq. However, there are few GSS tools specifically designed for scATAC-seq, and the applicability and performance of RNA-seq GSS tools on scATAC-seq data remain to be investigated. We systematically benchmarked ten GSS tools, including four bulk RNA-seq tools, five single-cell RNA-seq (scRNA-seq) tools, and one scATAC-seq method. First, using matched scATAC-seq and scRNA-seq datasets, we find that the performance of GSS tools on scATAC-seq data is comparable to that on scRNA-seq, suggesting their applicability to scATAC-seq. Then the performance of different GSS tools were extensively evaluated using up to ten scATAC-seq datasets. Moreover, we evaluated the impact of gene activity conversion, dropout imputation, and gene set collections on the results of GSS. Results show that dropout imputation can significantly promote the performance of almost all GSS tools, while the impact of gene activity conversion methods or gene set collections on GSS performance is more GSS tool or dataset dependent. Finally, we provided practical guidelines for choosing appropriate pre-processing methods and GSS tools in different scenarios.

## Introduction

Assay for Transposase-Accessible Chromatin using sequencing (ATAC-seq) is a powerful and the most widely used epigenomic technique for interrogating chromatin accessibility on a genome-wide scale [1]. In particular, the advent of single-cell ATAC-seq (scATAC-seq) has made it possible to profile chromatin-accessibility variations in single cells, which allows to illuminate chromatin-based gene regulation with an unprecedented cellular resolution and discover new cell subpopulations [2, 3]. One of the ultimate goals for analyzing single-cell chromatin accessibility data is to quantitatively understand the relationship between the variation of chromatin accessibility and that of the expression of nearby genes [4]. A first step toward this goal is to link regulatory DNA elements with their target genes on a genome-wide scale and predict gene activity (GA) score by modelling the chromatin accessibility at the gene level. Several tools are currently in progress to convert chromatin accessibility signals to GA scores, including Cicero [4], MAESTRO [5], ArchR [6], snapATAC [7], and Signac [8]. The inferred GA scores facilitate the integrative analysis of single-cell RNA-seq (scRNA-seq) and scATAC-seq data, and the scores of key marker genes can be used for accurate annotation of cell types as if in scRNA-seq [4, 6, 9].

In addition to single gene analysis, gene set analysis, analogue to pathway analysis, has become a routine step for analyzing gene expression data, which has proven to be effective in estimating the activity of pathways or transcription factors (TFs) for uncovering transcriptional heterogeneity and disease subtypes [10–12]. In single-cell RNA-seq studies, gene set scoring (GSS), or commonly referred to as pathway activity transformation, has been broadly conducted to quantify the enrichment and relevance of gene sets in individual cells. GSS converts the gene-level data into gene set-level information; gene sets contain genes representing distinct biological processes (e.g., the same Gene Ontology annotation) or pathways (e.g., the Molecular Signature Database (MSigDB) [13]). Therefore, GSS helps to decipher single-cell heterogeneity and cell-type-specific variability by incorporating prior knowledge from functional gene sets or pathways [14, 15]. A wide spectrum of GSS tools have been designed for scRNA-seq data, such as Pagoda2 [16], Vision [17], and AUCell [18], which infer pathway-level information from the gene expression profile for the characterization of transcriptional heterogeneity of cell populations. Similarly, gene sets with dynamic regulatory potentials inferred from scATAC-seq can also be regarded as cell-type specific markers as if in scRNA-seq [5].

Single-cell ATAC-seq data and RNA-seq data have analogous characteristic structures, both of which suffer from similar sparsity and noise. In recent years, great breakthroughs have been made in the computational modelling of scRNA-seq data, such as dropout imputation, dimensionality reduction, cell type identification, GSS, and regulatory networks inference [19–22]. In contrast, the progress on computational modelling in the field of scATAC-seq lags far behind that of scRNA-seq [23, 24]. As a compromise, many scRNA-seq analysis methods are directly applied to scATAC-seq data. For example, Liu et al. [25] benchmarked tools dedicated to imputing scRNA-seq data (e.g., MAGIC [26] and SAVER [27]) for recovering dropout peaks in scATAC-seq data and found that most scRNA-seq imputation tools can be readily applied to scATAC-seq data. Tools for alignment, quality control, peak calling, and differential peak analysis for RNA-seq and/or ChIP-seq data are widely used for ATAC-seq data [23]. This series of evidence indicates that GSS tools for scRNA-seq could in principle be applicable to scATAC-seq as well. However, due to the close-to-binary nature and extreme sparsity of the scATAC-seq data, it remains elusive whether these limitations would distort or confound the results produced by the direct application of RNA-seq methods to scATAC-seq. To the best of our knowledge, currently only one tool, UniPath [28], provides a function dedicated to scoring gene sets for scATAC-seq, therefore, it is timely and imperative to further investigate the applicability and performance of more GSS tools designed for bulk or single-cell RNA-seq on scATAC-seq data.

Currently the performance of GSS tools designed for bulk or single-cell RNA-seq on scRNA-seq data sequenced with diverse scRNA-seq protocols has been comprehensively evaluated. Zhang et al. [15] evaluated the performance of eleven pathway activity transformation tools on 32 scRNA-seq datasets and found Pagoda2 [16] exhibited the best overall performance. Holland et al. [14] compared the performance of six TFs or pathway activity estimators on simulated and real scRNA-seq data, which found that bulk tools can be applied to scRNA-seq, partially outperforming scRNA-seq tools. These studies focused only on scRNA-seq, to the best of our knowledge, there has been no systematic benchmark study to evaluate the performance of GSS tools on scATAC-seq data. Here we systematically evaluated the performance of ten GSS tools using ten scATAC-seq datasets, including four tools designed for bulk RNA-seq, five tools designed for scRNA-seq, and one method proposed for scATAC-seq. The performance was quantitively evaluated under four scenarios of dimensionality reduction, clustering, classification, and cell type determination, which are critical steps of single-cell analysis in most scRNA-seq and scATAC-seq studies. Our benchmark results provide abundant evidence that GSS tools designed for RNA-seq are also applicable to scATAC-seq. Using three matched scATAC-seq and scRNA-seq datasets, results showed that the performance of GSS tools for scATAC-seq data on clustering cells or distinguishing cell types was comparable to that for scRNA-seq. In particular, the performance of several GSS tools designed for RNA-seq exceeds the current only method dedicated to scATAC-seq, under diverse evaluation scenarios. Moreover, we evaluated the impact of data preprocessing of scATAC-seq on GSS, including dropout imputation and GA transformation. Benchmark results show that dropout imputation can significantly promote the performance of almost all GSS tools. In contrast, the performance of different GA transformation methods varies greatly across different GSS tools and different datasets. In addition, we also evaluated the performance of GSS tools using different gene set collections in the context of clustering and found that different GSS tools and different datasets have different degrees of robustness to different gene collections. Our benchmark results provide practical guidelines for choosing appropriate GSS tools for raw scATAC-seq data or data after dropout imputation, and also provide important clues on how to preprocess the scATAC-seq data for more effective GSS.

## Results

### Overview of the benchmark workflow

We benchmarked ten GSS tools, including four tools for bulk RNA-seq (PLAGE [29], z-score [30], ssGSEA [31], and GSVA [32]), five tools for scRNA-seq (AUCell [18], Pagoda2 [16], Vision [17], VAM [33], and UniPath [28]) and one function provided in the UniPath for scoring gene sets from scATAC-seq (hereinafter called UniPathATAC), using ten real scATAC-seq datasets with different number of cells and cell types (Figure 1). UniPathATAC can score gene sets directly from scATAC-seq data, using the peak-cell matrix as the input to obtain the gene set score matrix. In contrast, the input of RNA-seq GSS tools is the gene-cell matrix, thus the peak-level profile obtained from scATAC-seq data needs to be converted into the GA matrix, using a GA transformation tool. Four GA tools, including MAESTRO [5], Signac [8], ArchR [6], and snapATAC [7], were examined. MAESTRO obtains the GA matrix from the peak-cell matrix, while other three GA tools from the fragment file (Materials and methods). Unless otherwise specified, Signac was used as the default GA conversion tool as it runs fast and has good performance in our preliminary test. But we also conducted in-depth evaluation on the impact of different GA tools on GSS. Moreover, the pipeline for evaluating GSS tools involves an additional preprocessing step -- imputation of dropout peaks. We adopted three popular imputation tools developed for scRNA-seq (MAGIC [26], DrImpute [34], and SAVER [27]) and one tool designed for scATAC-seq (SCALE [35]). It should be noted that the imputation is performed on the peak-cell matrix rather than the fragment file, therefore, only MAESTRO [5] can be used for GA conversion from the imputed data. In addition, we examined six gene set collections from MSigDB (version 7.1), including KEGG (Kyoto Encyclopedia of Genes and Genomes), GO:BP (GO Biological Process), GO:MF (GO Molecular Function), GO:CC (GO Cellular Component), REACTOME, and TFT (Transcription Factor Target) (Table S1). Unless otherwise specified, KEGG that contains 186 gene sets in MSigDB was used as the default prior information. We benchmarked GSS tools under diverse scenarios of dimensionality reduction, clustering, classification and cell type determination. Each GSS tool was used to obtain the gene set score matrix from each scATAC-seq dataset (hereafter called GSS-ATAC), which was then evaluated in the context of each evaluation scenario.

**Figure 1.**
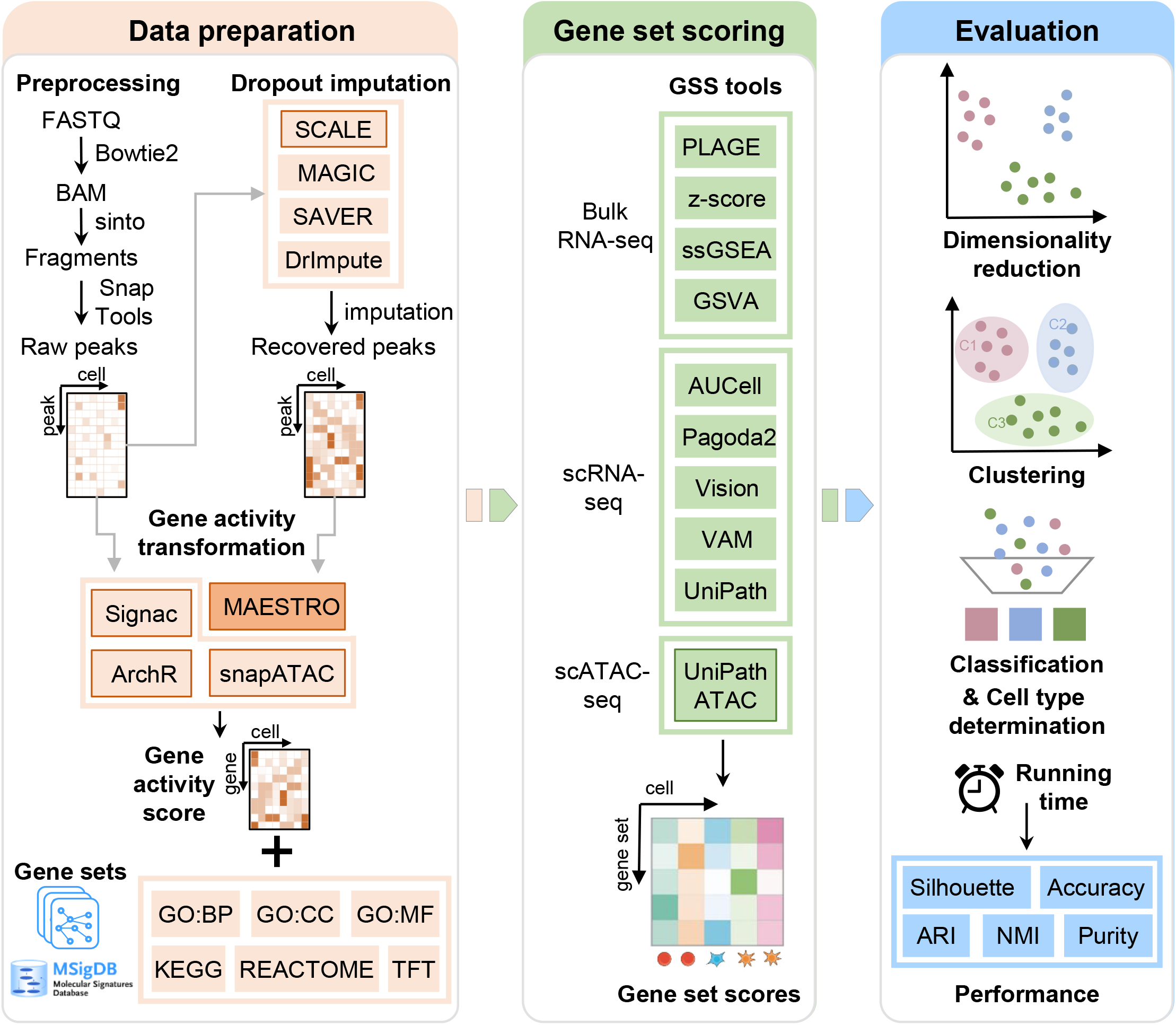
Overview of the benchmark workflow. Before applying GSS tools, scATAC-seq dropout peaks can be recovered by imputation tools and then the peak-level open-chromatin profile is converted into gene-level activity scores using GA transforming tools. Using gene sets from MSigDB as prior information, ten GSS tools are benchmarked in the context of diverse evaluation scenarios of dimensionality reduction, clustering, classification and cell type determination based on a variety of performance indicators. Tools marked with solid borders, including SCALE, the four GA tools and UniPathATAC, are specifically designed for scATAC-seq. MAESTRO can be used for GA transformation from both raw peaks and recovered peaks, while other three GA tools can be only applied to raw peaks as they require a fragment file which is not available for the imputed peak data. GSS, gene set scoring; scATAC-seq, single-cell assay for transposase accessible chromatin using sequencing; GA, gene activity; MSigDB, the molecular signatures database.

### GSS tools designed for RNA-seq are applicable to scATAC-seq

We used three matched datasets of scATAC-seq and scRNA-seq that are derived from the same cells, including Brain, PBMC3K, and PBMC10K (Table S2), to examine whether GSS tools designed for RNA-seq are applicable to scATAC-seq data. First, we used Signac to convert the peak-cell matrix to the GA matrix and then performed each GSS tool to obtain the GSS-ATAC matrix. We also used the nine RNA-seq GSS tools to score gene sets for the matched scRNA-seq data to obtain the corresponding gene set score matrix for scRNA-seq (hereafter called GSS-RNAseq). Then the performances of different GSS tools were evaluated by dimensionality reduction measured by Silhouette, clustering measured by ARI and classification measured by accuracy based on the GSS-ATAC or the GSS-RNAseq matrix obtained by different tools. We conducted the pipeline for each dataset and then calculated the average value of each performance indicator of the three datasets. For both scRNA-seq and scATAC-seq data, two methods, Pagoda2 and PLAGE, generally provide better performance than other methods in terms of all the three performance indicators (Figure 2A). Other GSS tools exhibit comparable and moderate performance. Although the performance of GSS tools on scRNA-seq and scATAC-seq is comparable, most GSS tools provide slightly better performance on scRNA-seq than on scATAC-seq. This is not unexpected because that these tools, except for UniPathATAC, were designed for RNA-seq and the reference cell types of scATAC-seq datasets were determined by the scRNA-seq data rather than scATAC-seq. Still, the consistency between clustering results obtained by GSS-ATAC and the reference cell types, measured by ARI, is even slightly higher than that of GSS-RNAseq obtained by several tools, including GSVA, VAM, and Vision (GSVA: 0.51 *vs*. 0.47; VAM: 0.38 *vs*. 0.36; Vision: 0.50 *vs*. 0.49). In particular, the performance of the two tools with the best performance, Pagoda2 and PLAGE, is higher than UniPathATAC, a method designed specifically for scATAC-seq, under all evaluation schemes (e.g., ARI of Pagoda2 = 0.60, PLAGE = 0.57, UniPathATAC = 0.55). Moreover, 2D embeddings of both GSS-ATAC and GSS-RNAseq matrices obtained by different GSS tools show comparable discrimination of the cell types (Figure 2B).

**Figure 2.**
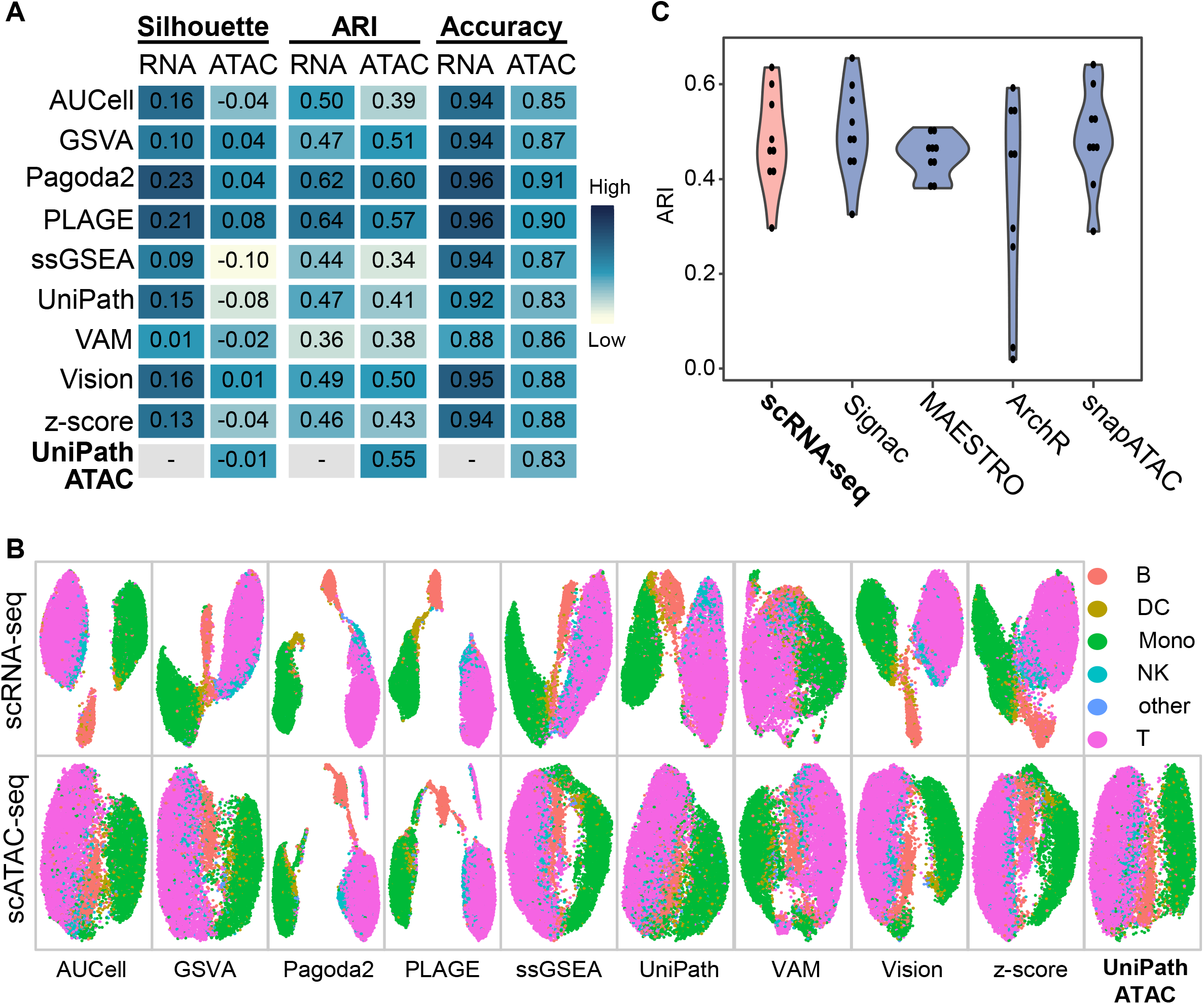
GSS results using matched datasets of scATAC-seq and scRNA-seq. **A.** Comparison of the performance of GSS tools on scRNA-seq (RNA) and scATAC-seq (ATAC) data in the context of dimensionality reduction measured by Silhouette, clustering measured by ARI, and classification measured by accuracy. Signac was employed to convert the peak-cell matrix into the gene-cell activity matrix, and KEGG gene sets were used as prior information. Three datasets including Brain, PBMC3K, and PBMC10K were used and the average performance was calculated. **B.** UMAP visualization of cell types using gene set scores obtained by applying different GSS tools on scRNA-seq and scATAC-seq PBMC10K data, respectively. The plot was created using the DimPlot function provided in the Seurat package. **C.** Comparison of the impact of different GA transformation tools on GSS of the PBMC10K data. Signac, MAESTRO, ArchR, and snapATAC were used for transformation and then ten GSS tools were applied on the GA matrix for scoring gene sets. Each violin plot summarizes ARI scores of the ten GSS tools, with each dot representing one tool. *P* values of Wilcoxon Rank Sum test used to compare ARI values between the scRNA-seq group and the other four groups of Signac, MAESTRO, ArchR, and snapATAC are 0.60, 0.60, 0.22, and 0.86, respectively. ARI, adjust random index; UMAP, uniform manifold approximation and projection.

In addition to Signac, we also used three other GA tools for transforming the peak profile to the gene-level activity scores and then calculated the GSS-ATAC matrix using different GSS tools. Results on the PBMC10K data show that the GSS-ATAC matrix based on the GA matrix obtained by different GA tools yields comparable ARI score to that using scRNA-seq data (Figure 2C), demonstrating again the applicability of RNA-seq GSS tools to scATAC-seq. Among the four GA tools, ArchR is less robust than other three GA tools for the PBMC10K data (Figure 2C). Taken together, these results preliminarily show that GSS tools designed for RNA-seq have comparable performance on both scRNA-seq and scATAC-seq data and thus are applicable to scATAC-seq data. In the following benchmark evaluation, we used more scATAC-seq datasets and considered different factors, including pre-processing steps, gene set collections, and GA methods, to evaluate different GSS tools more comprehensively.

### Evaluation of GSS tools using different scATAC-seq datasets

Having preliminarily demonstrated that GSS tools designed for RNA-seq are applicable to scATAC-seq, next we used eight scATAC-seq datasets (Table S2), which are from human and mouse with number of cells ranging from 500 to 10K, to further evaluate the performance of different GSS tools. Generally, the performance of GSS tools is highly dependent on datasets (Figure 3). Regardless of the evaluation scenario, the performance of all tools on Hematopoiesis, Leukemia, and SNAREmix is extremely poor, significantly lower than that on other five datasets. We then examined the raw scATAC-seq data to check whether the datasets with generally poor GSS results have low data quality. Indeed, we found significantly lower consistency between the clusters and the reference cell types of the three datasets with poor GSS results than the other five datasets (Figure S1). Although different GSS tools have varied performance on different datasets, Pagoda2 and PLAGE perform overall better than other tools. For example, the average ARI scores of all the eight datasets of Pagoda2 and PLAGE are much higher than that of the third tool UniPathATAC (Pagoda2 = 0.32, PLAGE = 0.30, UniPathATAC = 0.24). Of note, UniPathATAC is specially designed for scATAC-seq. Similarly, according to the scenario of classification, the average accuracy of PLAGE and Pagoda2 is also much higher than other tools (PLAGE=0.72, Pagoda2=0.67, other tools=0.62). These results revealed that the performance of the scATAC-seq specific tool, UniPathATAC, is only moderate, which is generally lower than that of two GSS tools for RNA-seq, Pagoda2 and PLAGE, suggesting again the feasibility of applying RNA-seq GSS tools to scATAC-seq data.

**Figure 3.**
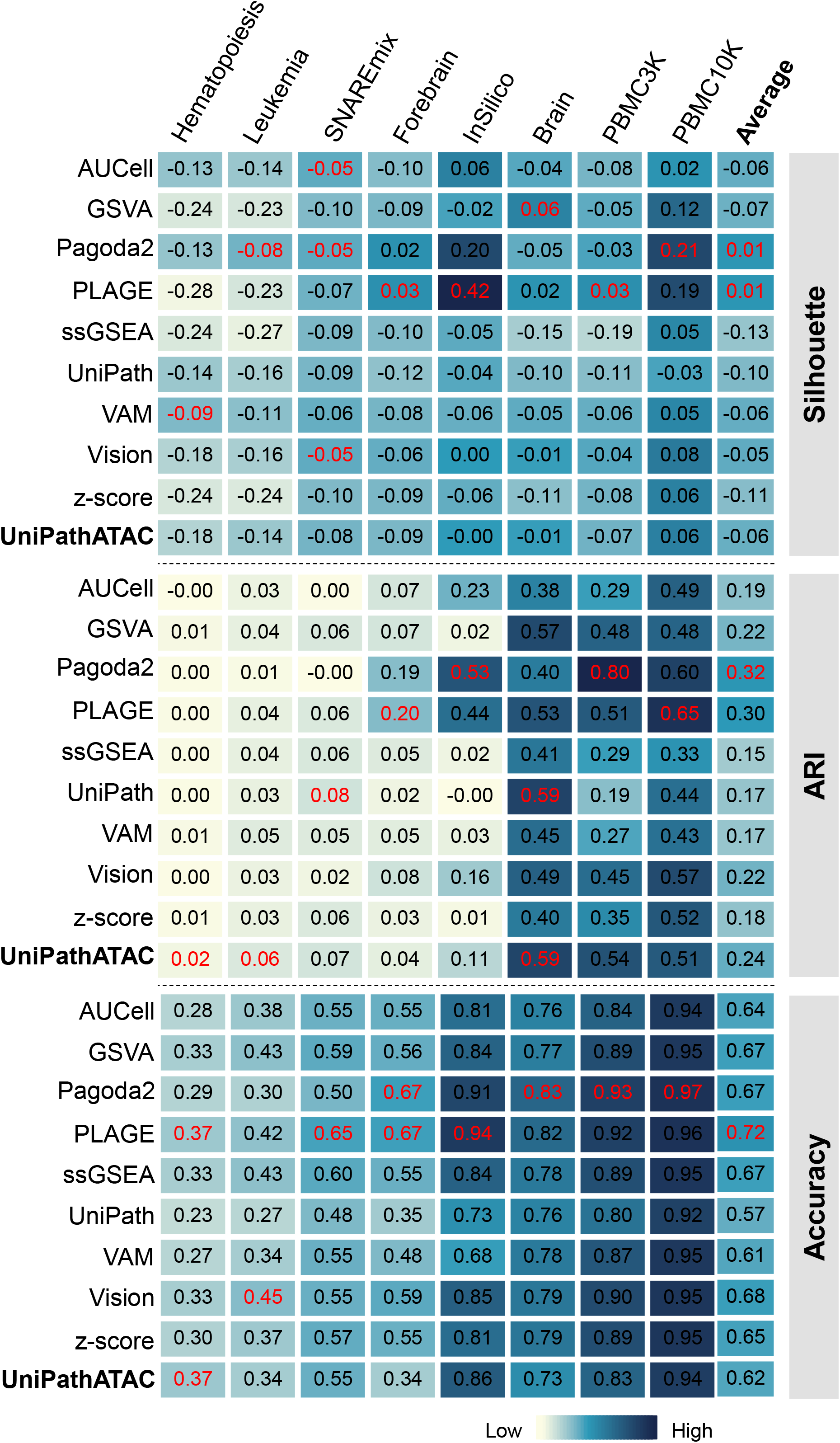
Comparison of the performance of GSS tools. The comparison was performed in the context of dimensionality reduction measured by Silhouette, clustering measured by ARI and classification measured by accuracy. In each column, the index values of the top performer for the respective dataset are displayed in red. The ‘Average’ column is the average score of each row.

### Evaluation of the impact of dropout imputation on GSS

Similar to scRNA-seq, scATAC-seq is plagued by extremely high sparsity and noise, therefore single-cell dropout peaks are usually recovered before downstream analysis. In contrast to the considerable progress that has been made in dropout imputation of scRNA-seq data, much fewer imputation tools for scATAC-seq are available. Till now, SCALE [35] is the only imputation method specially designed for scATAC-seq. A previous benchmark study [25] suggested that imputation tools designed for scRNA-seq are also applicable to scATAC-seq. Therefore, in addition to SCALE, we also considered three widely used scRNA-seq imputation tools, including MAGIC, DrImpute, and SAVER. Of note, the recovered peak-cell matrix can only be transformed into gene-cell activity matrix by MAESTRO, whereas the other three GA tools cannot because they use the fragment file for GA conversion. The performance of different GSS tools was compared under three evaluation scenarios -- dimensional reduction, clustering, and classification, using nine scATAC-seq datasets.

In general, regardless of imputation methods or GSS tools used, the performance of GSS using recovered peak profile is significantly improved compared with that using the raw peak profile (Figure 4A). Among the four imputation methods, SCALE that is designed for scATAC-seq provides the overall best performance, ranking first or second in almost all comparisons. Among the three scRNA-seq imputation methods, the overall performance of DrImpute is the best, followed by MAGIC. Except that the performance of SAVER is apparently the worst in most cases, the performance of the other three tools is relatively close. Moreover, the impact of the same imputation tool on the performance of different GSS tools is quite consistent, and no GSS tool relies on a specific imputation method. Next, we examined in detail the change of ARI scores of different GSS tools before and after imputation by SCALE under the clustering scenario (Figure 4B). In almost all cases, regardless of datasets or GSS tools, ARI scores based on recovered data are increased significantly. However, the performance improvement of different datasets after imputation varies greatly; the increase of ARI value under Leukemia, Hematopoiesis, and Brain is much slighter than that under other six datasets. Moreover, after imputation, the performance of different GSS tools on the same dataset also varies greatly. For example, after imputation, the ARI score of different tools on InSilico varies from 0.41 by UniPathATAC to 0.88 by Vision; the ARI score on GM12878HL varies from 0.02 by VAM to 0.79 by Pagoda2. In addition, the performance ranking of these tools changes after imputation. Pagoda2 and PLAGE are top performers using the raw data (Figures 2 & 3), while their ranking falls to a medium level after imputation. The performance of almost all tools has been greatly improved using data after imputation, but none is obviously the best -- several tools, including GSVA, Vision, Pagoda2, ssGSEA, and AUCell, achieve comparably good performance. Interestingly, the ARI score of Pagoda2 on raw data of InSilico and PBMC3K is much higher than that of other tools, however, the performance after imputation is even lower than that before imputation or most other tools. This result indicates that the impact of data imputation for a tool that already performs well on the raw data may be limited. In contrast, some GSS tools have very poor performance before imputation, while a substantial improvement was obtained after imputation. For example, the ARI score of Vision on the InSilico raw data is only 0.13, while it is increased greatly to 0.88 using data after imputation. The UMAP visualization of the GSS-ATAC matrix obtained from the InSilico data shows significantly more distinguishable cell types using data after imputation (Figure 4C). These results demonstrate that the performance of GSS tools can be significantly improved by the incorporation of the imputation step in data preprocessing, particularly for those GSS tools having poor performance on the raw data.

**Figure 4.**
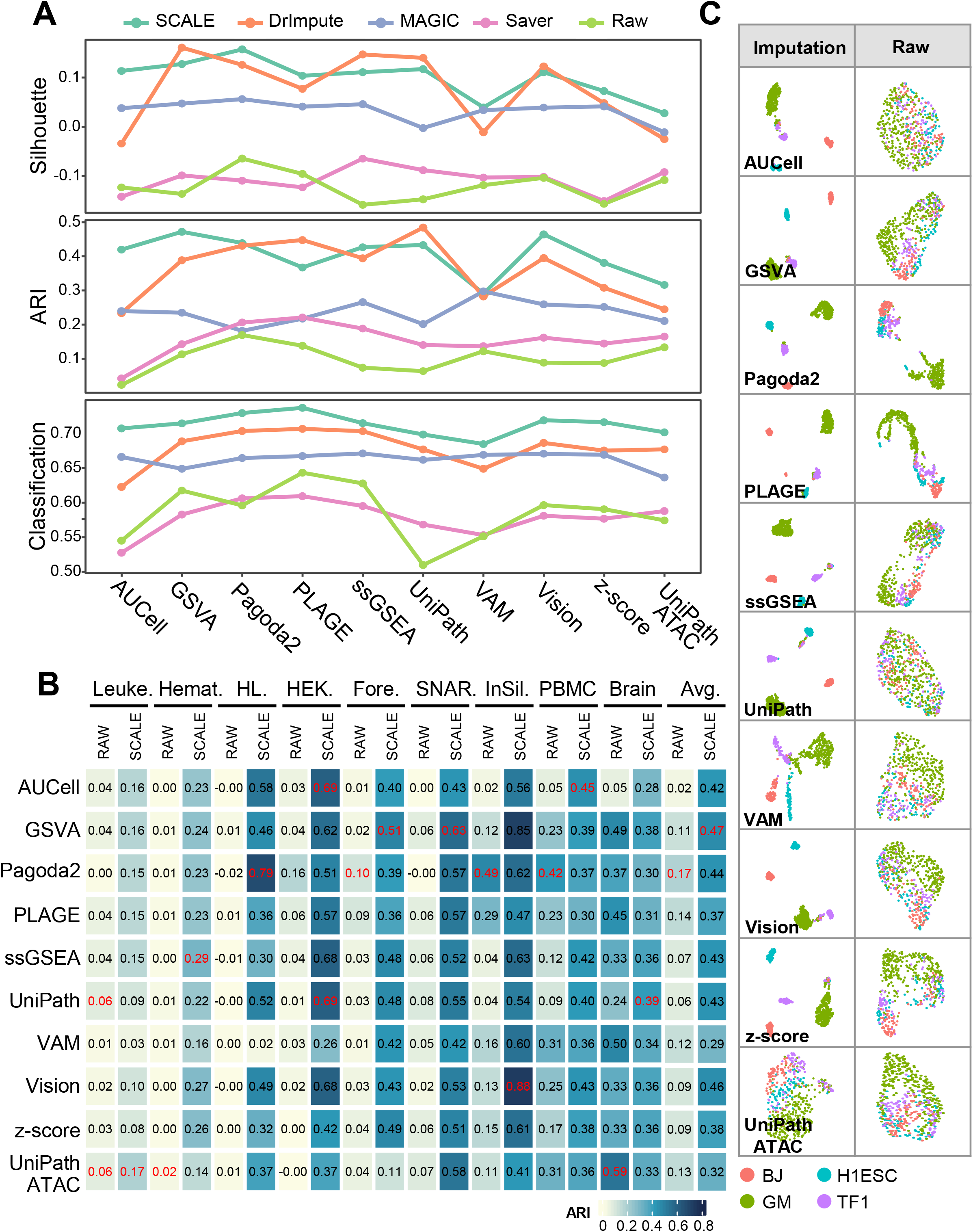
Comparison of the impact of different dropout imputation tools on GSS. **A.** Average performance of GSS tools on nine scATAC-seq datasets before or after imputation in the context of dimensionality reduction measured by Silhouette, clustering measured by ARI and classification measured by accuracy. **B.** The change of ARI scores of different GSS tools before and after imputation by SCALE. In each column, the index value of the best performer for the respective dataset is coloured in red. The ‘Avg.’ column is the average score of each GSS tools on the nine datasets before (RAW) or after imputation (SCALE). **C.** UMAP visualization of cell types using gene set scores obtained from the raw or imputed peak profile of the InSilico data by each GSS tool. Datasets: Leuke., Leukemia; Hemat., Hematopoiesis; HL., GM12878HL; HEK., GM12878HEK; Fore., Forebrain; SNAR., SNAREmix; InSil., InSilico; PBMC., PBMC3K.

### Evaluation of GSS tools by the enrichment analysis of marker gene sets

Next, we used marker genes of known cell types as the reference to further evaluate the accuracy of cell type recognition using gene sets quantified by different GSS tools (Materials and Methods, Table S3). Considering the abundance of cell types and the availability of cell marker information in the CellMarker database [36], here we used the two PBMC datasets with 25 sub-types for evaluation. ssGSEA has the highest accuracy of cell type recognition when only the top one to three gene sets were used (Figure 5). For example, when identifying cell types only based on the top one gene set, the accuracy of ssGSEA is ~71%, which is much higher than other tools (Vision = 51% in the second place). Several other tools also achieve comparable accuracy to ssGSEA when using ≥ 3 top gene sets, including VAM, Pagoda2, and Vision, which reach an accuracy of > 82% using top five gene sets. Surprisingly, for PLAGE which has comparable performance with Pagoda2 in other evaluation scenarios (Figures 2 & 3), none of the top gene sets identified by PLAGE is enriched on correct cell types. In particular, although UniPathATAC is designed purposely for scATAC-seq, its performance is consistently lower than several other GSS tools for RNA-seq. Taken together, among the ten GSS tools, six tools, including ssGSEA, VAM, Pagoda2, Vision, AUCell, and z-score, provide overall better performance than other tools. UniPathATAC and GSVA rank at the second level, while UniPath and PLAGE perform the worst.

**Figure 5.**
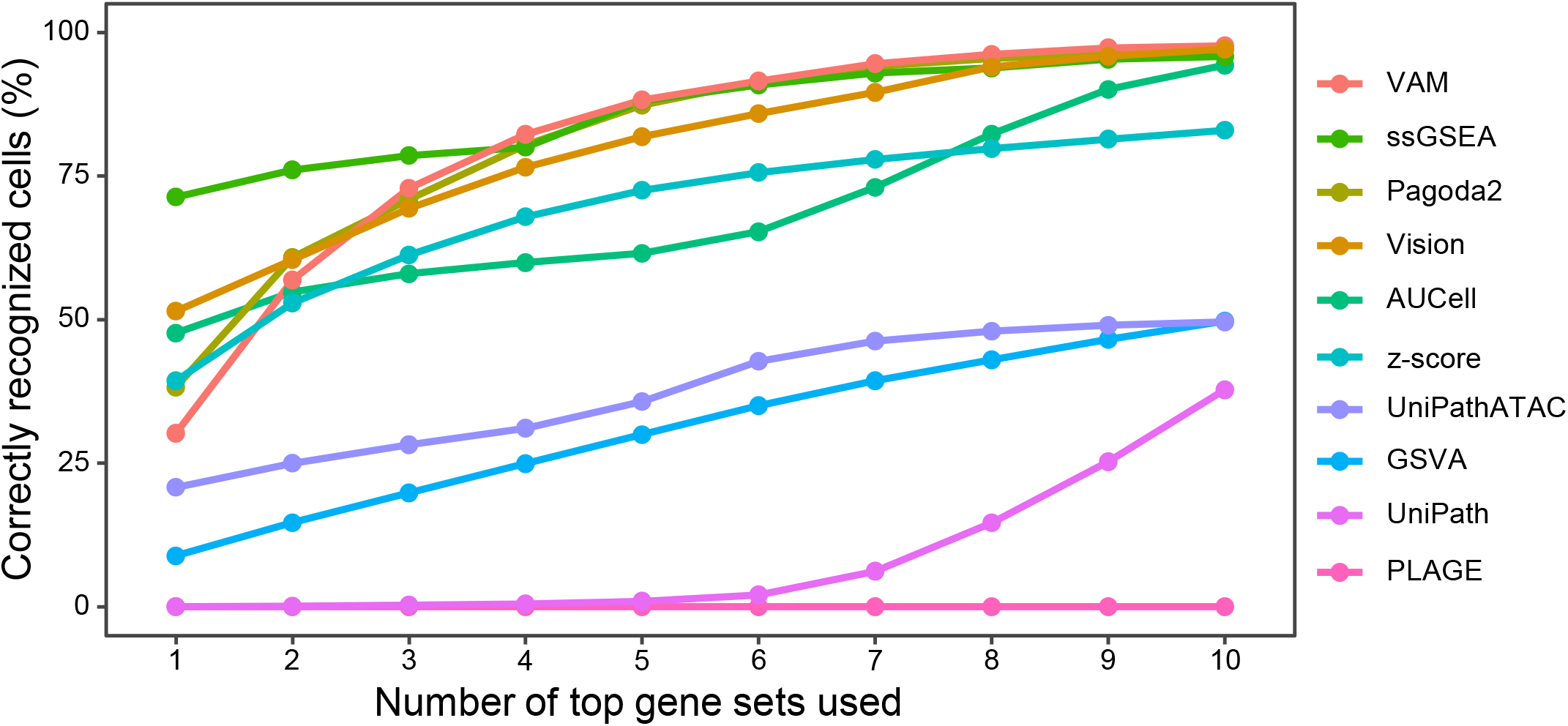
Evaluation of the enrichment and relevance of gene sets in single cells quantified by different GSS tools. PBMC3K and PBMC10K datasets were used, with six main cell types and 25 sub-types. Marker genes of 467 known cell types from the CellMarker database were used as the reference. Each GSS tool was used to score the 467 marker gene sets for each PBMC dataset, and the top *N* gene sets ranking by the gene set score can be obtained for each cell. If a cell’s cell type falls within cell types of the top *N* gene sets, then the cell is considered as correctly recognized. The Y-axis denotes the average percentage of cells annotated with correct cell type of the two PBMC datasets based on the results of each GSS tool. The X-axis denotes the number of top gene sets used for cell type recognition.

### Evaluation of the impact of GA transformation on GSS

GA conversion is a necessary step before using RNA-seq GSS tools on scATAC-seq data. Here we evaluated the performance of different GA tools by calculating the correlation between the GA profile from scATAC-seq and the gene expression profile from scRNA-seq, using three matched scRNA-seq and scATAC-seq datasets (Brain, PBMC3K, and PBMC10K). Generally, Signac and snapATAC provide better consistency between GA inferred from scATAC-seq and gene expression level from scRNA-seq than MAESTRO and ArchR (Figure 6A). Using the SCALE-imputed data for GA conversion by MAESTRO, the consistency measured by correlation is increased (*P* value < 5.8e-108 between MAESTRO/SCALE and MAESTRO/raw for each dataset), suggesting that imputation could increase the performance of GA conversion. Next, we compared the effect of GA tools on GSS using more scATAC-seq datasets. Since GA tools except for MAESTRO are only applicable to the raw scATAC-seq peak profile, we used the raw data without imputation for evaluation. GA matrix obtained by GA tools were used as the input for the ten GSS tools to score gene sets. There is no clear consensus on which approach is the best; no GA method has significantly higher impact on the performance of all GSS tools than other methods (Figure 6B). Among the ten GSS tools, the performance variation of different GA methods on AUCell and UniPath is greater than that on other GSS tools. Among the four GA tools, the performance of different GSS tools on GA matrix obtained by Signac and snapATAC is more robust and relatively higher than that by MAESTRO or ArchR. Moreover, different from other GSS tools for RNA-seq that can only score gene sets from the GA matrix, UniPathATAC can score gene sets directly from the peak profile without GA transformation, while its performance is inferior than several GSS tools designed for RNA-seq, such as Pagoda2 and PLAGE. Collectively, Signac and snapATAC provide relatively better results than MAESTRO and ArchR in both evaluation scenarios, whereas MAESTRO has the unique ability to obtain GA from imputed data.

**Figure 6.**
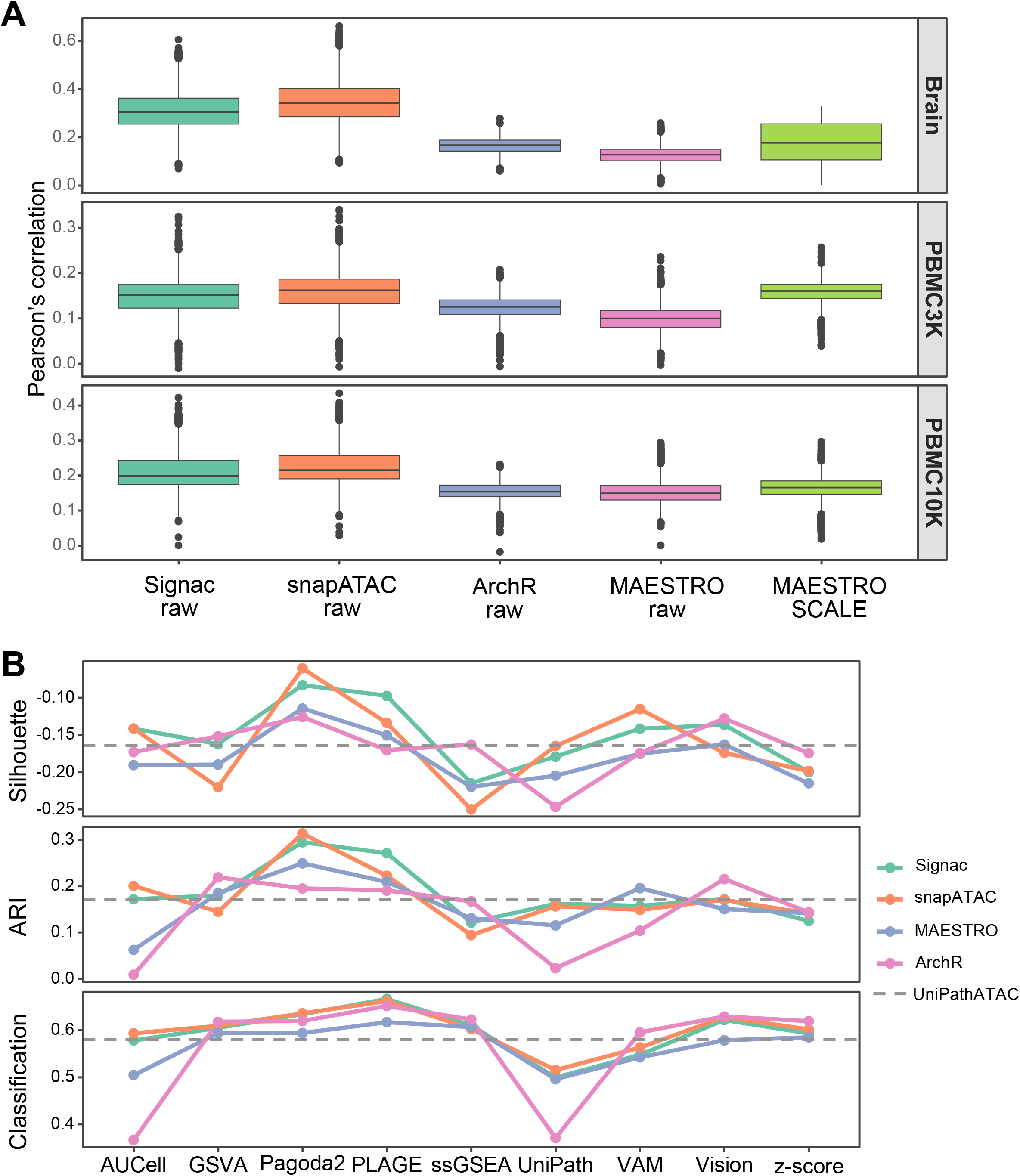
Comparison of the impact of different GA transformation tools on GSS. **A.** The correlation between the GA profile obtained by four GA transformation tools from scATAC-seq and the gene expression profile from scRNA-seq for three matched scRNA-seq and scATAC-seq datasets. Labels with ‘raw’ means the GA tools was performed on the raw scATAC-seq profile, while ‘SCALE’ means that MAESTRO was used on the SCALE-imputed scATAC-seq profile. B. Average performance on seven datasets in the context of dimensionality reduction measured by Silhouette, clustering measured by ARI and classification measured by accuracy. Results of UniPathATAC that is designed for scATAC-seq without needing GA transformation are displayed as horizontal dotted lines for comparison. For three of the ten scATAC-seq datasets used in this study (GM12878HEK, GM12878HL, and SNAREmix), the fragment file that is needed for GA conversion of Signac, snapATAC, and ArchR was not available, therefore, the remaining seven datasets were used here for evaluation, including Leukemia, Hematopoiesis, Forebrain, InSilico, PBMC3K, PBMC10K, and Brain.

### Evaluation of the impact of different gene set collections on GSS

Next, we investigated the impact of six gene set collections from MSigDB (Table S1) on the performance of GSS tools, using nine scATAC-seq datasets. In the evaluation pipeline, we used SCALE for dropout imputation, followed by MAESTRO for GA transformation. Then we applied different GSS tools to each GA matrix to calculate the GSS-ATAC matrix based on each gene set collections, and evaluated the performance in the context of clustering. The impact of different gene set collections on GSS performance is not as evident as that of imputation tools (Figure 7A *vs*. Figure 4A). The average ARI score using TFT or GO:BP is slightly lower than that using other four gene set collections (TFT = 0.367; GO:BP = 0.389; others: 0.4 to 0.419). Moreover, different GSS tools have different degree of robustness to different gene set collections on different datasets (Figure 7B). For four datasets (Brain, Hematopoiesis, Leukemia, and PBMC3K), the performance of all GSS tools is relatively stable, regardless of which gene set collection is used (Figure 7C). In contrast, for the other five datasets, the performance of different GSS tools is more affected by gene set collections. For example, for InSilico which shows overall high performance, AUCell, GSVA, and Vision are much less sensitive to gene sets than other tools (Figure 7B). Among the ten GSS tools, the performance of Vision and UniPath is the least affected by gene sets, while UniPathATAC is the most sensitive to gene sets (Figure 7C). In particularly, Pagoda2 is the top performer on raw scATAC-seq data according to our evaluation (Figures 2A & 3), however, its robustness to different gene sets is only moderate (Figures 7B & C). Overall, Vision has relatively more robust and generally high performance across different gene set collections.

**Figure 7.**
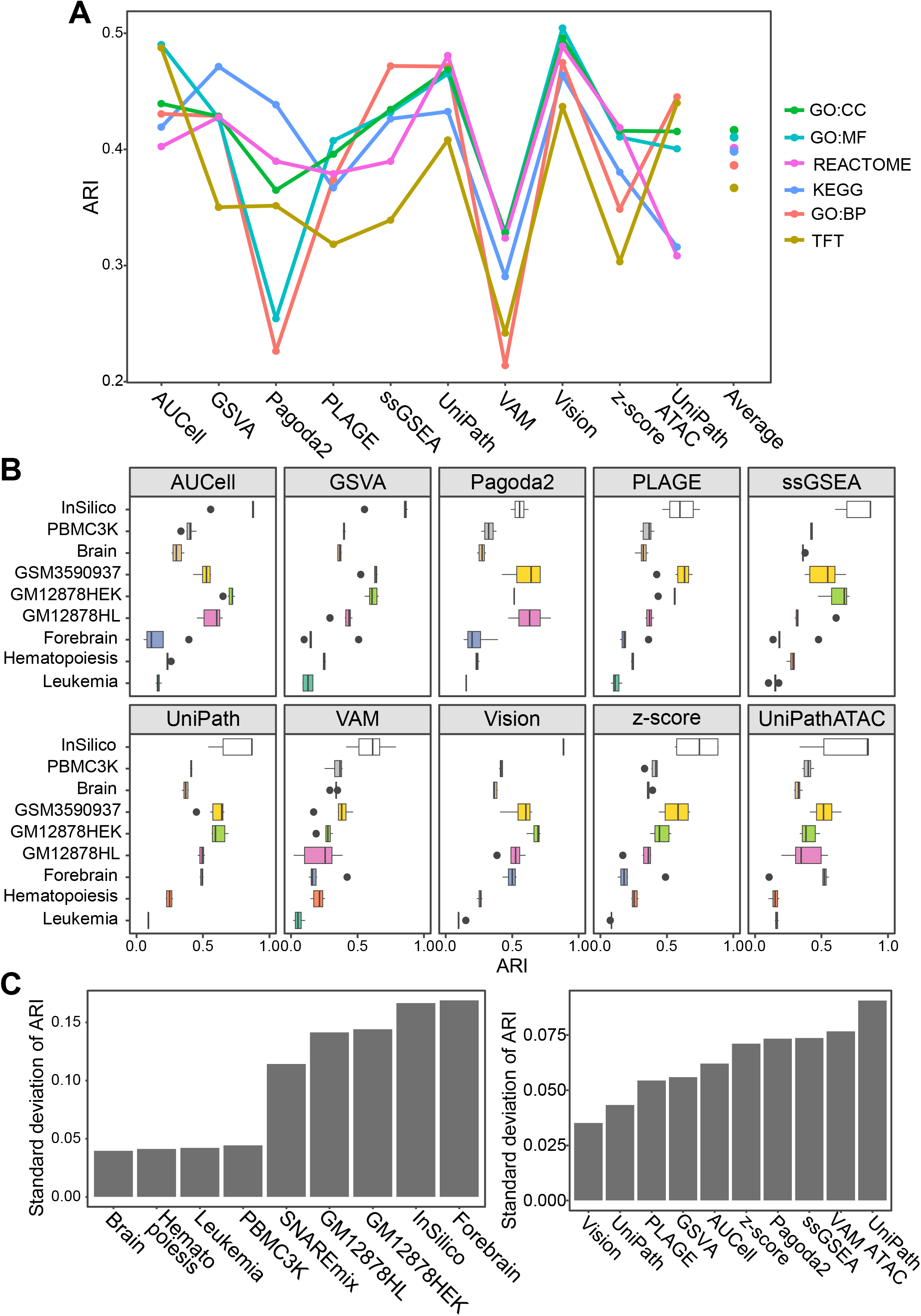
Comparison of the impact of different gene sets on GSS. **A.** Average ARI score of ten GSS tools on nine scATAC-seq datasets using six gene set collections from MSigDB. Dots in the ‘Average’ column represent the average ARI score of all GSS tools using the respective gene set collection. Average ARI scores: GO:CC = 0.419; GO:MF = 0.412; REACTOME = 0.401; KEGG = 0.4; GO:BP = 0.388; TFT = 0.368. Dropout peaks in each scATAC-seq dataset were recovered by SCALE, followed by MAESTRO for GA transformation. **B.** Each boxplot summarizes the ARI scores by applying a GSS tool on the six gene set collections. KEGG, Kyoto encyclopedia of genes and genomes; GO, gene ontology; GO:BP, GO biological process; GO:MF, GO molecular function; GO:CC, GO cellular component; TFT, transcription factor target. **C.** Standard deviation (SD) of ARI scores on different datasets (left) or GSS tools (right). To obtain the SD for each dataset, the average of the SD of ARI scores of all GSS tools using different gene set collections was calculated. To obtain the SD for each GSS tool, SD of ARI scores of the GSS tool on each dataset using different gene set collections was calculated. Then the average of SD on different datasets for each GSS tool was calculated.

### Running time evaluation

The computing speed of Vision and z-score is significantly faster than that of other tools. Even when the number of cells and gene sets increases, the running time only increases slightly (Figure 8). In contrast, GSVA and VAM run fast when the data size is small, while the running time increases significantly with the increase of data size. Among these tools, ssGSEA and UniPath take significantly more computing time than other tools. Nevertheless, among these experiments, it only takes up to six hours (Unipath: 328.84 min) even for the longest case by these two tools. PLAGE and Pagoda2, which show the best performance on the raw data, are quite efficient, which are second in line to the fastest tools, Vision and z-score. However, Pagoda2 failed to complete calculation in some cases, which needs to be used with caution. According to the calculation speed, the ranking for the top three tools with overall high performance on data after imputation is Vision > Pagoda2 > ssGSEA. In addition, UniPathATAC, a tool specially designed for scATAC-seq, has a medium computing speed, which is close to Pagoda2.

**Figure 8.**
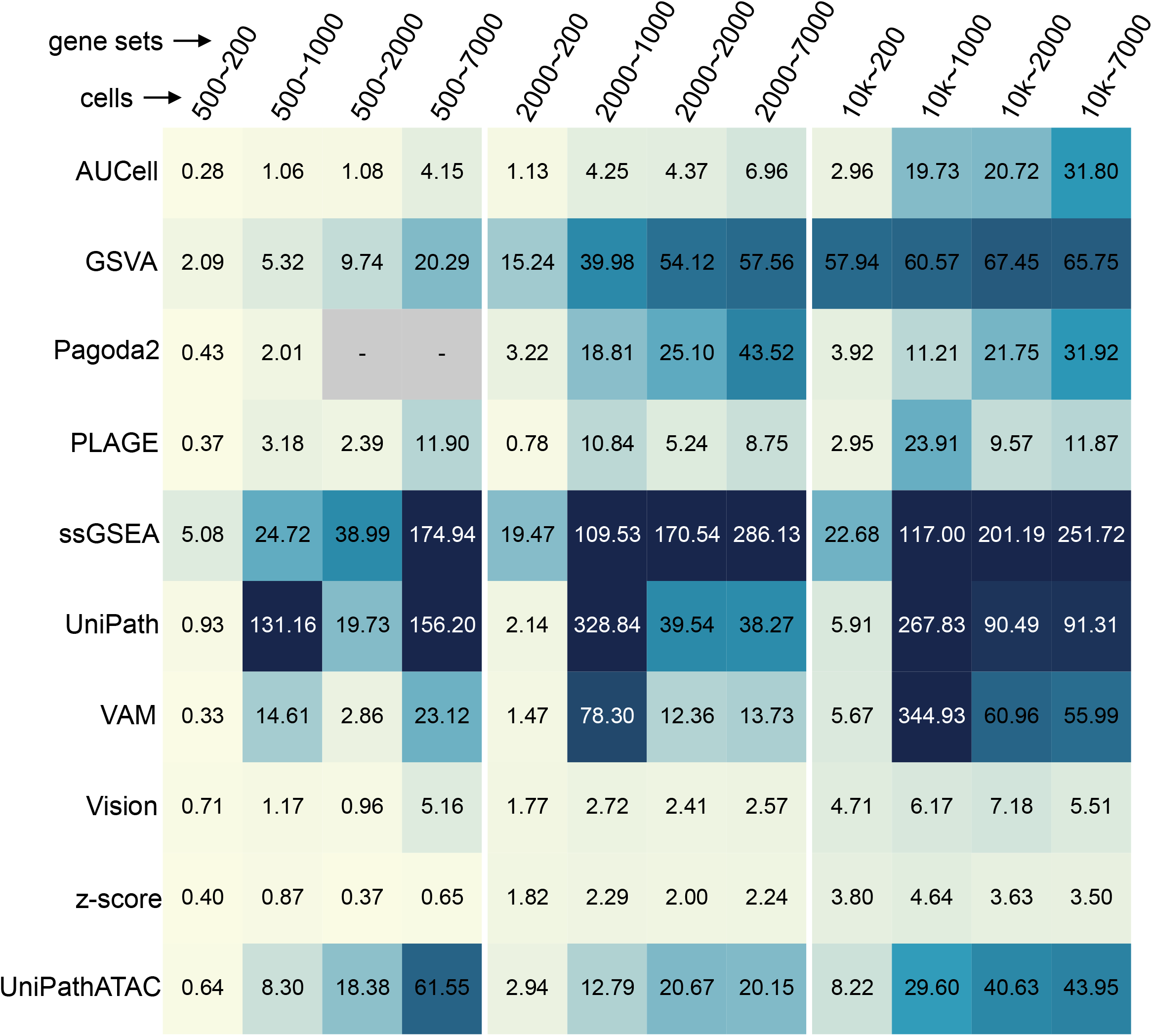
Evaluation of running time (in minute) of different GSS tools. Three datasets were tested, including InSilico, Hematopoiesis and PBMC10K, which contain approximately 500, 2000, and 10,000 cells, respectively. Four gene set collections were used, including KEGG, TFT, REACTOME, and GO:BP, which contain approximately 200, 1000, 2000, and 7000 pathways, respectively. Cases where Pagoda2 failed to complete the calculation are marked with ‘-’.

### Practical guidelines for choosing GSS tools

Here we summarized the performance of different GSS tools on ten scATAC-seq datasets in various evaluation pipelines in the context of clustering, considering different GA tools, imputation tools, and gene set collections (Figure 9A). For the preprocessing of scATAC-seq data in the GSS pipeline, our results showed that dropout imputation can significantly improve the GSS results, and SCALE or DrImpute provide overall better performance than the other two imputation tools. In contrast, using different GA tools or gene set collections has much less impact on GSS results. Regardless of gene set collections, for peak-cell data after dropout imputation by SCALE (only MAESTRO can be used for GA transformation in this case), Vision and GSVA show an overall better performance on the SCALE-recovered data than other GSS tools (average ARI: GSVA = 0.47, Vision = 0.46, others = 0.29 to 0.44). For raw peak-cell data, Pagoda2 in conjunction with snapATAC (ARI = 0.31) or Signac (ARI = 0.29) performs the best, followed by PLAGE. In particular, it is worth noting that RNA-seq GSS tools are only applicable to scATAC-seq when the peak-level open-chromatin profile of scATAC-seq has been converted into gene-level activity scores by GA tools. Although our benchmark demonstrates that dropout imputation greatly improves the performance of GSS tools, only MAESTRO can be applied to the recovered peak-cell matrix for GA transformation, while other GA tools cannot due to that the fragment file needed for GA conversion cannot be imputed.

**Figure 9.**
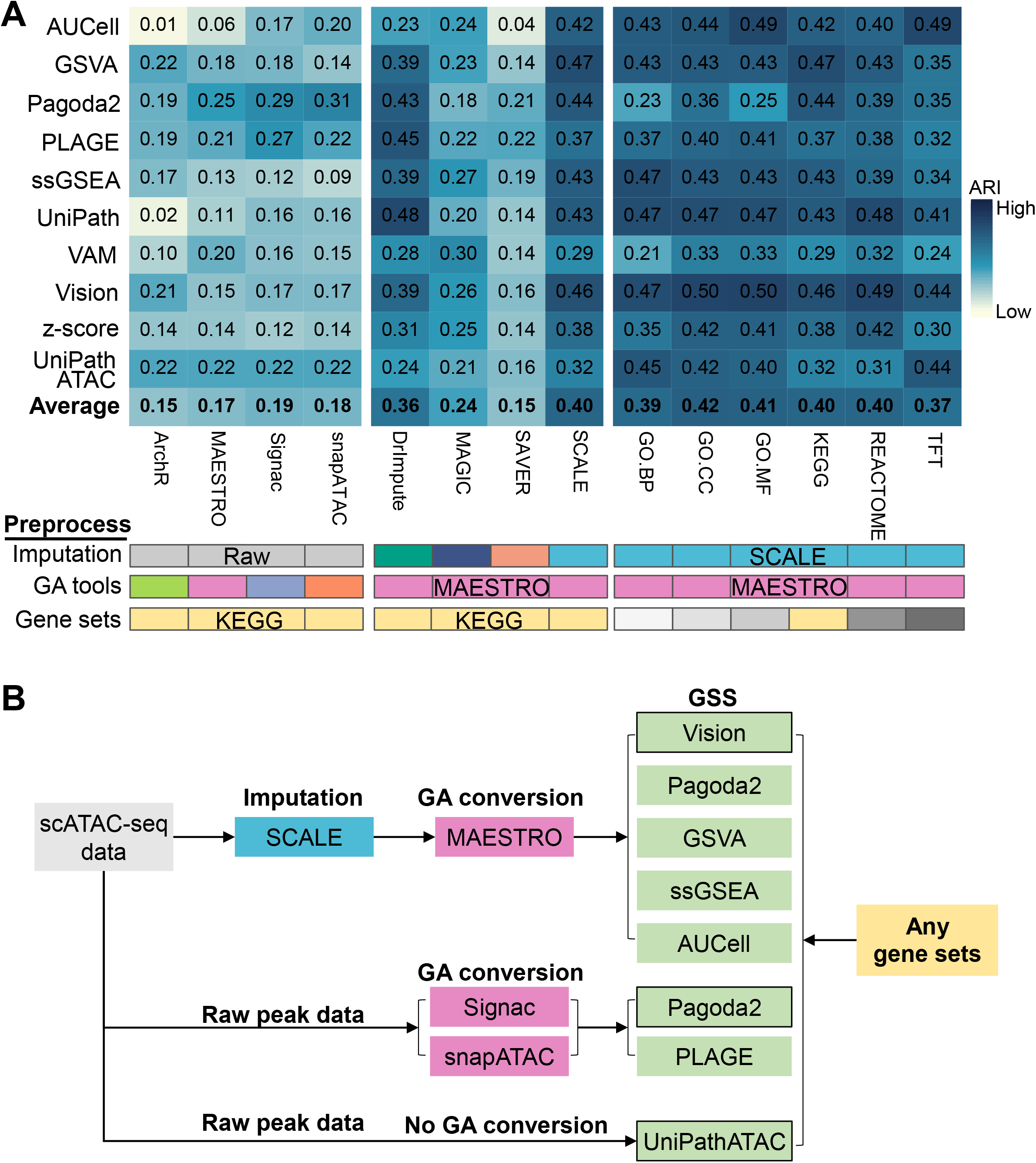
Summarization of the performance of different GSS tools in various evaluation pipelines measured by ARI. **A.** ARI scores of different scATAC-seq datasets were averaged. Cases guiding the tool recommendation are coloured in red. Each column denotes an evaluation task, which involves GA transformation with each of the four tools, dropout imputation (no imputation or imputation with each of the four tools), and selection of six gene set collections. Of note, when the dropout imputation is performed for the peak-cell matrix, only MAESTRO can be used for GA transformation because the other three GA tools are only applicable to the fragment file. **B.** Practical guidelines for choosing appropriate tools for GSS. The GSS tool with border is the most recommended tool with the best overall performance in the respective group.

Based on our comprehensive evaluation and unique features of different tools, we propose some practical guidelines for choosing appropriate tools for GSS (Figure 9B). For GSS from raw scATAC-seq data without dropout imputation, we recommend two tools with overall best performance and high speed, PLAGE and Pagoda2, combined with snapATAC or Signac for GA transformation (Figures 2 and 8). Meanwhile, users can also use SCALE to recover the peak-cell profile, followed by GA conversion with MAESTRO, and then adopt Vision, which has relatively good performance (Figures 4, 5, and 7) and speed (Figure 8) for data after imputation. Since the performance of different GSS tools on data after imputation is greatly improved and becomes closer (Figure 4), users can also try multiple GSS tools with comparable performance to Vision, such as GSVA, Pagoda2, ssGSEA, and AUCell, for comparative analysis, especially when the data size is small. If users want to perform GSS without GA conversion, then UniPathATAC is the only tool available at present. In addition, considering that different gene set collections have relatively limited and uncertain impact on the performance of GSS tools (Figure 7) but are important for biological interpretation, it is recommended to try different gene set collections in the GSS pipeline.

## Discussion

GSS has been widely conducted in bulk or single-cell RNA-seq studies, which helps to decipher single-cell heterogeneity and cell-type-specific variability by incorporating prior knowledge from functional gene sets or pathways. ScATAC-seq is a powerful epigenetic technique for interrogating single-cell chromatin-based gene regulation, and genes or gene sets with dynamic regulatory potentials can be regarded as cell-type specific markers as if in scRNA-seq. The GA score transformed from the chromatin accessibility profile of scATAC-seq is potentially a reliable predictor of gene expression and can be used for cell type annotation [4–8]. GA scores facilitate the use of RNA-seq GSS tools to score gene sets for scATAC-seq data. Taking the GSS results of the matched scRNA-seq datasets and those of UniPathATAC as the reference, we confirmed that RNA-seq GSS tools are applicable to scATAC-seq. First, we performed GSS for the matched scATAC-seq and scRNA-seq data from PBMCs and Brain, and found that the performance of GSS tools on scATAC-seq for clustering cells or distinguishing cell types was comparable to that on scRNA-seq (Figure 2). Second, by the enrichment analysis of marker gene sets for cell types using PBMC10K scATAC-seq data, we found that the top few (1-10) gene sets with high scores can be used to determine the cell types of most cells (Figure 5). Third, the comprehensive evaluation of various scATAC-seq datasets shows that several RNA-seq GSS tools, e.g., Pagoda2, PLAGE, and Vision, even have much better results under different evaluation scenarios than the GSS tool specially designed for scATAC-seq -- UniPathATAC (Figures 2–6). After demonstrating the applicability of RNA-seq GSS tools on scATAC-seq, we systematically evaluated 10 GSS tools and found that Pagoda2 and PLAGE have the best overall performance for the raw peak-cell profile, which is similar to the previous benchmark results of GSS tools on scRNA-seq data [15]. In particular, Pagoda2 is developed for scRNA-seq and PLAGE is for bulk RNA-seq, both of which are PCA-based RNA-seq methods but also provide good performance on scATAC-seq. Several previous studies have shown that GSS tools developed for bulk RNA-seq are applicable to scRNA-seq data [14, 15], and tools for scRNA-seq imputation is also widely used in recovering scATAC-seq dropouts [25]. Our benchmark further confirmed that GSS tools designed for RNA-seq is also suitable for scATAC-seq data.

We also comprehensively evaluated the impact of data preprocessing of scATAC-seq on GSS, including dropout imputation and GA transformation. We found that GSS results using data after imputation are significantly better than those using raw data, regardless of GSS tools or imputation tools (Figure 4). Among the four imputation tools, SCALE performs generally better than other three scRNA-seq tools, while the scRNA-seq tool DrImpute provides comparable performance to SCALE. Previously, Liu et al. [25] benchmarked multiple scRNA-seq imputation tools on scATAC-seq including MAGIC and SAVER, and found that MAGIC provides much better performance than SAVER. This is consistent to our observation that SAVER shows the worst performance on scATAC-seq data. Moreover, the two tools included in our benchmark that have overall high performance, SCALE and DrImpute, were not involved in the previous benchmark [25]. Particularly, the performance of Pagoda2 and PLAGE, which provide the best performance on raw data, is not significantly improved after imputation, while the performance of several other tools, including GSVA, Vision, Pagoda2, ssGSEA, and AUCell, is greatly improved after imputation, surpassing Pagoda2 and PLAGE (Figure 4). Compared to the positive impact of dropout imputation on GSS, the impact of different GA methods or gene set collections on GSS is uncertain and limited (Figures 6 & 7). Therefore, we recommend users to try different GA tools and different gene sets for GSS in practical applications. Moreover, we found that although the open-chromatin profile obtained from scATAC-seq data can be preprocessed using different imputation tools and different GA tools, GSS results are highly dependent on scATAC-seq datasets. Some datasets, such as Hematopoiesis and Leukemia, have extremely poor results regardless of the evaluation scenarios (dimensionality reduction, clustering or classification) or the representation of the data (peak profile, gene-level activity score or gene set score) (Figures 3,4, and S1). The low quality of the raw scATAC-seq data could be alleviated to some extent by dropout imputation rather than choosing a different GA tool. However, no matter how the raw data is preprocessed, GSS results on data with very poor quality of raw data often cannot reach the ideal level.

In our benchmark study, the performance of GSS tools was quantitively evaluated under four scenarios of dimensionality reduction, clustering, classification, and cell type determination. These scenarios, especially clustering, are critical steps of single-cell analysis in most scRNA-seq and scATAC-seq studies. We acknowledged that the ARI score that represents the consistency between the predicted cell type labels from clustering and the true reference is not high throughout our benchmarking of GSS tools (< 0.5 in most cases), which means that the clustering results solely based on gene set scores may be poor. However, for scATAC-seq data, which is even sparser than the already sparse scRNA-seq data, the ARI value is normally very low. For example, the ARI value in these pioneering scATAC-seq studies [25, 37–39] is also < 0.5 in most cases. Nevertheless, clustering is a routine step in most single-cell analysis pipelines and the outputs of different tools or methods are frequently used as the input for clustering algorithms to produce clustering results. Therefore, evaluating the clustering ability would be a useful measure for assessing the performance of different GSS tools. We estimated that the value of ARI can reflect the performance of different GSS tools under the clustering scenario. At the same time, the low ARI value indicates that the clustering results should be used in caution. Moreover, we also speculated that the low ARI value may be also due to the poor annotation or high similarity of some cell types, and/or the inability to completely restore the true cell types only through the scATAC-seq data. As such, integrating information of additional modalities with gene set scores, such as the gene expression profile from scRNA-seq and the peak-level profile from scATAC-seq, would help to obtain better clustering results for better cell type distinguishing.

Currently, matched scRNA-seq and scATAC-seq data on dynamic processes (e.g. differentiation of induced pluripotent stem cells) are increasingly available [40–44]. It would be interesting to examine whether and how well the cell transition trajectory could be inferred based on gene set scores obtained by different GSS tools. However, trajectory analysis is a more complex procedure that requires more biological interpretation than clustering analysis, and its results are difficult to quantify using performance indicators like ARI in clustering analysis. Nevertheless, evaluating GSS tools under the scenario of trajectory analysis could be a future direction upon the availability of appropriate quantification methods for evaluation the accuracy of trajectory inference.

## Material and methods

### Datasets

We used ten publicly available scATAC-seq datasets (Table S2), including InSilico [2], GM12878HEK [3], GM12878HL [3], Leukemia [45], Hematopoiesis [46], Forebrain [47], SNAREmix [48], and three matched datasets from 10X Genomics (Brain, PBMC3K, and PBMC10K) [8]. The InSilico dataset is an *in silico* mixture of four independent scATAC-seq experiments performed on different cell lines [2]. The GM12878HEK and GM12878HL datasets are mixtures of two commonly-used cell lines, respectively [3]. The Leukemia dataset includes mononuclear cells and lymphoid-primed pluripotent progenitor cells isolated from a healthy human donor, and leukemia stem cells and blast cells isolated from two patients with acute myeloid leukemia [45]. The Forebrain dataset is derived from P56 mouse forebrain cells [47]. The Hematopoiesis dataset was used to characterize the epigenome pattern and heterogeneity of human hematopoiesis [46]. The Brain, PBMC3K, and PBMC10K datasets are publicly available datasets generated by 10x Genomics [8], which jointly profiled mRNA abundance and DNA accessibility in human peripheral blood mononuclear cells (PBMCs) and human healthy brain tissue of cerebellum, respectively. The SNAREmix dataset is a mixture of cultured human BJ, H1, K562, and GM12878 cells [48]. These diverse datasets were generated from both microfluidics-based and cellular indexing platforms with substantially different number of cells and peaks, which were widely used in previous studies for benchmarking [25] or validating computational tools for scATAC-seq, such as scMVP [38], scABC [49], SCALE [50], and Signac [8]. We used Azimuth [51] to annotate cell types in the PBMC3K and PBMC10K datasets by label transfer from a publicly available multimodal PBMC reference dataset [51] and in Brain dataset by label transfer from the human cerebellum dataset [52]. Cell types of other datasets were obtained from relevant studies.

### Preprocessing of scATAC-seq data

For scATAC-seq datasets without publicly available peak-cell matrix, the raw FASTQ files downloaded from NCBI were aligned to the reference genome (human: hg19; mouse: mm10) using Bowtie 2 [53], resulting in alignment files of BAM format. Then these BAM files were used as inputs for MACS2 [54] for peak calling and then SnapTools (https://github.com/r3fang/SnapTools) was adopted to generate the peak-cell matrix. Similar to the previous study [55], we filtered peaks with read counts >=2 and present in at least 10 cells for InSilico, GM12878HEK and GM12878HL data. We filtered peaks with read counts >=2 and present in at least 50 cells for Forebrain. For Hematopoiesis, Leukemia, SNAREmix, Brain, PBMC3K and PBMC10K, we followed the routine preprocessing following the tutorial of Signac to filter peaks and cells.

We chose four tools for dropout imputation of scATAC-seq data, including SCALE [35] which is currently the only method specifically designed for scATAC-seq and three widely used scRNA-seq tools – MAGIC [26], DrImpute [34] and SAVER [27]. The peak-cell matrix was used as the input for these tools with default parameters for recovering dropout peaks. Of note, because Signac, ArchR, and snapATAC require a fragment file of the raw scRNA-seq data to calculate gene-level activity, we can only use MAESTRO [5] to obtain GA matrix directly from the recovered peak-cell matrix. We used liftOver [56] to convert coordinates between different genome versions, if necessary.

### GA conversion

The peak-level profile of scATAC-seq data needs to be converted into the gene-level activity before using RNA-seq GSS tools. We chose four GA tools, including MAESTRO [5], Signac [8], ArchR [6], and snapATAC [7], to transform the open-chromatin profile obtained from scATAC-seq into the gene-level activity scores. MAESTRO obtains a regulatory weight based on the distance from the peak center to the gene transcription start site, and associates it with the peak-cell matrix to get the gene activity score. Signac is used in the Seurat package [22] for GA conversion, which simply sums the gene body with the peaks that intersect in the 2-kbp upstream region in each cell. SnapATAC obtains a score for each gene by normalizing the number of fragments overlapping genes in cells. ArchR infers gene expression from chromatin accessibility by using a custom distance-weighted accessibility model. Among these tools, MAESTRO use the peak-cell matrix for GA conversion, while other three tools use the fragment file. The fragment file [8] is a coordinate-sorted file for storing scATAC-seq data, which contains five columns: chromosome, start coordinate, end coordinate, cell barcode, and duplicate count. This file can be generated from a BAM file using Cellranger or the Sinto package (https://pypi.org/project/sinto/). It should be noted that, only the peak-cell matrix rather than the fragment file can be imputed by imputation tools, therefore, only MAESTRO can be used for GA conversion on the peak-cell data after imputation.

We used three matched scRNA-seq and scATAC-seq datasets (Brain, PBMC3K, and PBMC10K) to evaluate the performance of different GA tools in predicting the gene expression level from scATAC-seq data. First, we used each GA tool to convert the raw peak-cell matrix into the GA matrix for each dataset. As MAESTRO is applicable to the imputed peak-cell profile, we also used MAESTRO to obtain the GA matrix based on the SCALE-imputed peak-cell matrix. Then we calculated the Pearson’s correlation between the raw or imputed GA profile from scATAC-seq and the gene expression profile from scRNA-seq for each cell. The correlation profiles of all cells obtained from the four GA tools for each matched scRNA-seq and scATAC-seq dataset were compared.

### GSS tools

Ten GSS tools were evaluated in our benchmark. We run these tools with default parameters according to the tutorials provided in the respective studies.

PLAGE (Pathway Level Analysis of Gene Expression) [29] scores gene sets for RNA-seq by singular value decomposition (SVD). The gene expression matrix is normalized, and the first coefficient of the right singular vector obtained by SVD is considered as the gene set score.

Combined z-score (z-score) [30] is a classic strategy to aggregate the expression of multiple genes. Gene expression is scaled by the mean and standard deviation of the cells. Then, gene expression levels of all genes within each gene set are averaged to score the gene set of each cell.

ssGSEA (Single Sample Gene Set Enrichment Analysis) [31] is an extension of GSEA. ssGSEA ranks genes by expression levels within each cell individually, then scores gene sets by enrichment analysis using random walk statistics such as Kolmogorov-Smirnov (K-S) statistic.

GSVA (Gene Set Variation Analysis) [32] utilizes the K-S statistic to assess gene set variation. GSVA first estimates the cumulative density function for each gene, using the classic maximum deviation method by default. The score matrix is obtained by calculating the score of the gene set from the gene density profile using the K-S statistic.

AUCell [18] employs the area under the curve (AUC) to calculate the enrichment of a pathway (i.e., gene set) in the expressed genes of each cell. AUCell first ranks genes based on their expression levels in each cell, resulting in a ranking matrix. The AUC of the recovery curve is then used to determine whether the gene set is enriched at top genes in each cell. To calculate AUC, only the top 5% of genes are used by default, which means to examine how many genes in the gene set are within the top 5% genes in the respective cell.

Pagoda2 (Pathway and Gene Set Overdispersion Analysis) [16] is a computational framework to detect cellular heterogeneity from scRNA-seq data. The method fits an error model to each cell to characterize its properties, and then renormalizes the residual variance for each gene in the cell. Then, the scoring matrix for each gene set is quantified by its first weighted principal component.

Vision [17] uses autocorrelation statistics to identify biological variation across cells, which performs directly on the manifold of cell-cell similarity. It first identifies the K-Nearest Neighbors (KNNs) of each cell to generate a KNN map of the cell, then the GSS matrix is calculated based on the average gene expression of each gene set.

VAM (Variance-Adjusted Mahalanobis) [33] is a fast and accurate method for cell-specific gene set evaluation, which is integrated with the Seurat framework to accommodate the characteristics of high technical noise, sparsity and large sample size of scRNA-seq data. It calculates cell-specific pathway scores to convert a gene-by-gene matrix into a pathway-by-pathway matrix that can be used for data visualization and statistical enrichment analysis.

UniPath [28] is a uniform approach for pathway and gene-set based analysis for both scRNA-seq and scATAC-seq. For scRNA-seq, it first converts gene expression profiles to p-values assuming a Gaussian distribution, according to the mean and variance of each cell. Then p-values of genes in each gene set are combined using Brown’s method and then an adjusted p-value is obtained for each gene set. For scATAC-seq, UniPath first highlights enhancers by normalizing read counts of scATAC-seq peaks using their global accessibility scores and performs a hypergeometric or binomial test using proximal genes of peaks, which then converts the open-chromatin profile to pathway enrichment scores for gene sets. UniPath provides functions for scoring gene sets in scRNA-seq and scATAC-seq, respectively. In this study, we referred to the method for scRNA-seq as UniPath and the method for scATAC-seq as UniPathATAC.

### Benchmarking gene set scoring tools

#### Cell type clustering

We evaluated the performance of different GSS tools in the context of unsupervised clustering, using Louvain which is imbedded in the Seurat package. Given a GSS-ATAC matrix obtained by a GSS tool, we employed PCA for dimensionality reduction and then performed Louvain clustering on the first 10 PCs. Louvain clustering provides a tuneable parameter ‘resolution’ for determining the number of clusters based on a binary search algorithm, which was set to 0.5 in our benchmark. We used ARI (Adjust Random Index), a widely-used indicator, to measure the consistency between two clustering results. The ARI is the adjusted value of the raw RI (Random Index) score; the RI computes a similarity metric between two clustering results by considering all sample pairs and counting pairs assigned in the same or different clusters in the predicted and true clusters (Eq. 1). An ARI close to 0 means random labelling and ARI = 1 means perfect matching of the two clustering results. ARI is calculated with the ‘adjustedRandIndex’ function in the mclust [57] package.

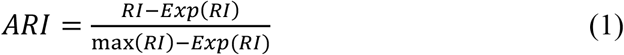

#### Dimensionality reduction

We first performed dimensionality reduction by PCA on the GSS-ATAC matrix obtained by a GSS tool with Seurat (PCs = 10). Then UMAP (Uniform Manifold Approximation and Projection) [58] was performed with the first 10 PCs and the average Silhouette width of all cells was calculated using the ‘silhouette’ function provided in the R package cluster. The Silhouette score was used to evaluate the performance of dimensionality reduction for each GSS-ATAC matrix. Silhouette score ranges from −1 to 1, with a high value indicating that cells of the same cell type group together and are far from cells of a different type. The silhouette score for cell *i* is defined as:

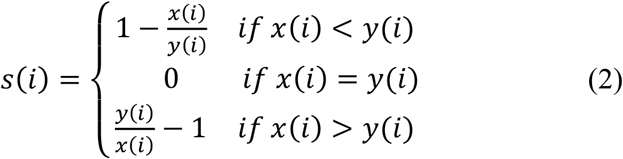

Here, *x*(*i*) and *y*(*i*) is the average distance from cell *i* to all other cells in cell *i*’s cluster and cell *i*’s nearest cluster, respectively.

#### Classification

To evaluate the performance of GSS tools in the context of classification, we implemented a multi-normal logistic regression model with k-fold cross-validation using the Python scikit-learn package. The inverse of the regularization strength of the multinormal logistic regression model was set to 1. The parameter *k* of the k-fold cross-validation was set to 5. Gene set scores in the GSS-ATAC matrix were scaled between 0 and 1 before model training and testing. The classification accuracy of the test dataset is calculated.

#### Enrichment analysis of marker gene sets

Similar to the previous study [28], we used marker genes of known cell types as the reference to examine whether gene sets scored by different GSS tools are enriched on known cell types. We obtained human marker genes from CellMarker [36] to make a collection of gene sets for 467 cell types (Table S3) and then organized these gene sets as the form of the gene set representation in MSigDB. Each GSS tool was used to score these marker gene sets for each scATAC-seq dataset to obtain a GSS-ATAC matrix. Based on the GSS-ATAC matrix, for each cell the top *N* gene sets ranking by the gene set score can be obtained. If a cell’s cell type falls within cell types of the top *N* gene sets, then the cell is considered as correctly recognized. Finally, given a scATAC-seq dataset, the percentage of cells annotated with correct cell type was calculated for each GSS tools.

#### Running time evaluation

We used scATAC-seq datasets and gene sets with different sizes to test the running time of GSS tools. Three datasets with different orders of magnitude were used for evaluation, including InSilico, Hematopoiesis and PBMC10K, which contain approximately 500, 2000 and 10K cells, respectively. Four sources of gene sets with different sizes were selected from MSigDB, including KEGG (186 pathways), TFT (1133 pathways), REACTOME (1797 pathways) and GO:BP (7350 pathways). The computer processor for evaluation is intel@Xeon(R) CPU E5-2680 v4 @ 2.40GHz × 56. One CPU core is allocated to each task of running a GSS tool on a dataset with given gene sets. Only the running time of the GSS tool is counted, excluding the time consumption of data and package loading, preprocessing, data imputation and gene activity conversion.

## CRediT author statement

**Xi Wang**: Investigation, Methodology, Data curation, Formal analysis. **Qiwei Lian**: Investigation, Methodology, Data curation, Formal analysis. **Haoyu Dong**: Data curation. **Shuo Xu**: Data curation. **Yaru Su**: Formal analysis. **Xiaohui Wu**: Conceptualization, Writing - original draft, Writing - review & editing, Supervision, Project administration, Funding acquisition. All authors read and approved the final manuscript.

## Acknowledgements

This work was supported by the National Natural Science Foundation of China (Grant No. T2222007 to XW). Funding for open access charge: National Natural Science Foundation of China.

## Supplementary materials

**Figure S1.**
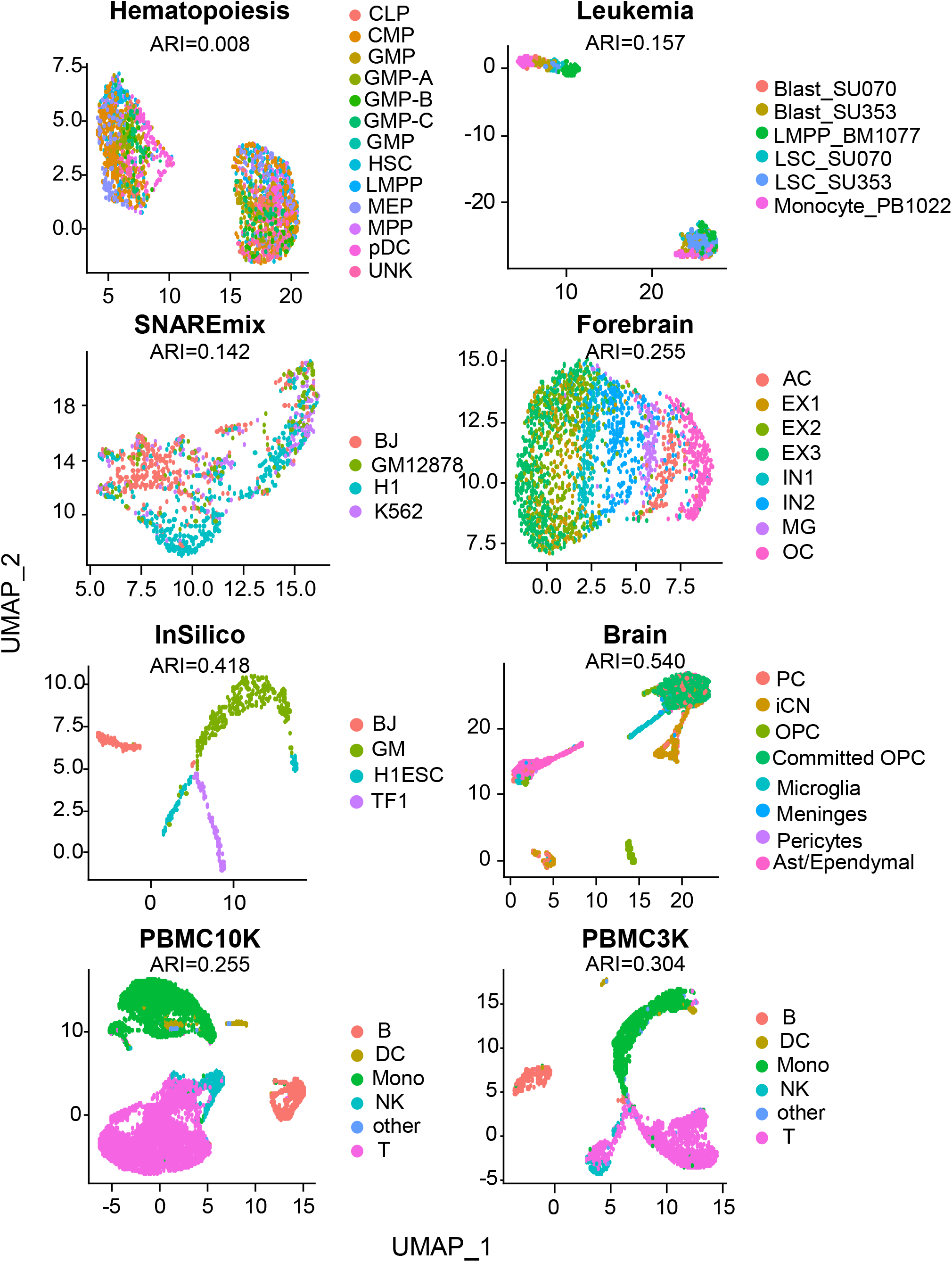
UMAP plots showing 2D-embeddings of the raw peak-cell matrix of eight scATAC-seq datasets.

**Table S1 Size of gene set collections used in this study**

**Table S2 Detailed information of scATAC-seq and scRNA-seq datasets used in this study**

**Table S3. Human marker gene sets collected from the CellMarker database**

